# Proteomic study for the prediction of μCT imaging with iodine

**DOI:** 10.1101/2025.04.22.649994

**Authors:** Valentin Wesp, Lisa-Marie Barf, Heiko Stark

## Abstract

Iodine-based staining techniques are commonly used in histological imaging and micro-computed tomography (µCT) due to iodine’s affinity for binding to specific molecules. However, the basis for tissue-specific contrast has not yet been sufficiently explored. In this study, we analyse the human proteome at four different levels: individual proteins, protein families, tissues with additional expression values for selected proteins, and organs as a distinct combination of different tissues. At each level, we try to identify proteins/groups with high potential for iodine binding, especially those rich in aromatic heterocyclic amino acids. Using bioinformatic methods, we evaluate the occurrence of aromatic/non-aromatic heterocyclic, carbocyclic, and the remaining 15 amino acids in 20,650 proteins, 1,487 families, 57 tissues, and 16 organs. At the protein level, structural proteins such as titin, nebulin, obscurin, mucin, filaggrin and hornerin have a high absolute number of aromatic heterocyclic amino acids, which could explain the high µCT contrast in muscle, skin and mucosal tissues. At the next level, however, structural families (such as the *Laminin*-family) rank significantly lower in comparison. These results are reflected in tissues and organs for which protein expression is available. Here, no significant correlations between the enrichment of heterocycles and the intensity of iodine staining can be observed. Furthermore, the enrichment of amino acids in each tissue/organ is relatively similar and shows no significant difference. Our results provide a general basis for iodine-based tissue imaging and serve as a potential starting point for future research, e.g. for cross-species applications and for the structural and functional effects of iodination.

## Introduction

Iodine has been used in histology for many years to stain tissue. The most common and cheapest form is Lugol’s solution, which is a dissolved mixture of iodine and potassium iodide. It was first described by Jean Guillaume Auguste Lugol in the early 1800s^1^. In solution, iodine (I2) and potassium iodide (KI) additionally form triiodides (*I*3^+^) and potassium ions (*K*^*−*^)^2^. This solution causes the normal, non-cornified squamous epithelium of the normal oesophagus, for example, to discolour differently from pathologically altered tissue^1,3^. This is explained by the affinity of glycogen for iodine-containing substances^4,5^. Lipid staining is also known^6,7^. The staining mechanism for high-contrast 3d imaging is not yet fully understood^5^.

It is known that pure iodine is an inert halogen that usually requires catalysis to form a bond^8^. These reactions include oxidations of disulfide bridges (Cys), hydrophilic and ion-ion interactions (amino & carbonyl groups). Another possibility of staining can be explained by intercalation processes into helical structures of starch with triiodide, leading to an intense blue-black colour^9,10^. Electron-rich aromatic heterocycles (pyrrole, furan, or thiophene), on the other hand, can react very easily directly with iodine^11^. Such aromatic heterocyclic compounds are a widespread class in nature, with important roles in biological processes. They include the nucleic bases of RNA and DNA, cytochromes, cobalamins, some natural colourants and odorants, as well as the amino acids histidine and tryptophan^12,13^. Proline, being a saturated five-membered heterocycle, does not follow Hückel’s rules and is therefore not aromatic. As a result, it does not react spontaneously with pure iodine.

Two reviews by Metscher^5,14^ show the strength of iodine staining. He compares several contrast agents (Gallocyanin-chromalum, I2E, I2M, IKI, PTA, Osmium tetroxide) for the staining of soft tissue. Iodine provides results comparable to those of the other contrast agents. However, it is preferable in terms of usability and toxicity^14^. It is also shown that iodine primarily stains yolk, skin, eyes, nerves, and muscle tissue, which are protein-rich tissues.

There is also a special analysis method, called the iodine value determination, that measures the degree of unsaturated compounds in fats^15^. This utilises the fact that interhalogens (iodine chloride) specifically add to olefinic bonds and can be used for quality control. However, it should be noted that the iodine has to be bonded; otherwise, it will not attach to the alkenes. This method also explains why iodine can be used to stain lipids.

So why is iodine used to stain soft tissue for µCT examinations? This is because soft tissue has a very homogeneous density and offers little contrast. Iodine helps here because of its atomic radius. The radius of a covalently bonded iodine atom is around 133 picometres, which falls within the wavelength range of X-rays: 10 to 10,000 picometres^16^. X-rays are therefore easily absorbed by the iodine atoms and, due to the fact that iodine rarely occurs in animal tissue, are suitable for imaging iodine-rich areas.

From the above-mentioned facts, aromatic heterocycles and olefinic double bonds can be selectively stained and spatially visualised through imaging techniques. This raises the question of which metabolites or proteins are being labelled. In this study, we were particularly interested in identifying tissues rich in aromatic heterocycles (histidine and tryptophan) and, therefore, expected to exhibit intense staining.

## Methods

For our study, we retrieved the human proteome (containing 20,650 proteins) in FASTA format from the [UniProt database](https://www.uniprot.org/) (accessed 14 February 2025) using a custom Python script (v3.12). We chose a human dataset as a proof-of-concept due to its high level of curation and relevance. After filtering for uncurated sequences, 322 were removed, leaving 20,328 curated sequences in our dataset. The remaining sequences were then analysed for their amino acid composition, quantified as absolute counts as well as proportions of aromatic residues (histidine and tryptophan), non-aromatic heterocyclic residues (proline), carbocyclic residues (phenylalanine and tyrosine), and all other canonical amino acids.

To compare specific staining in µCT, intensities and expression values for the aforementioned proteome were downloaded from three selected databases: the [MorphoSource Database](https://www.morphosource.org/) (MS), the [Human Protein Atlas](https://www.proteinatlas.org/) (HPA) and the [ProteomicsDB](https://www.proteomicsdb.org/proteomicsdb/) (PtDB).

For the MS database, this includes iodine-stained volume data for five mammalian species (000039697, 000039783, 000042252, 000042257, 000371199)^17^. Differentiating tissues was not possible due to the low resolution compared to normal histological staining. Therefore, the volume data were restricted to organs. The analysis was performed using [3D Slicer](https://www.slicer.org/) to obtain normalised intensity values for each available organ.

Next, we grouped proteins into four levels to obtain distinct enrichment values for aromatic heterocycles: protein, family, tissue, and organ levels. For the protein level, enrichments were simply calculated as the relative proportions of aromatic heterocycles in each protein. For the family level, proteins were grouped based on the [HUGO Gene Nomenclature Committee](https://www.overleaf.com/project/67ffab336790b5278883e8be) (HGNC) database, which contains 1,447 families. Of the 20,328 proteins in the dataset, 4,760 could not be assigned to any family. At the protein level, the relative proportions of aromatic heterocycles in each family were determined.

For the tissue and organ level, expression values from the HPA and PtDB databases were included in the analysis. The HPA database consists of two datasets, transcriptomics data that provide a snapshot of gene expression in an adult human, and proteomics data that quantify protein levels using specific antibodies via immunohistochemistry. However, the HPA proteomics dataset does not contain any actual values; it only contains expression levels labelled ‘Not detected’, ‘Low’, ‘Medium’, and ‘High’. To simplify things and enable comparison with other datasets, the labels were mapped to the values ‘0’, ‘1’, ‘2’, and ‘3’, in that order. The last dataset is proteomics data from the PtDB, which quantifies protein abundance using mass spectrometry. Proteins with an expression value of zero across all tissues and organs were discarded from the respective dataset. It is important to note that expression data were not available for each protein in all three databases.

The tissue and organ classifications of the proteins was based on [Terminologia Anatomica IDs](http://terminologia-anatomica.org/en/) (TA98), which comprised 57 tissues and 16 organs. These IDs are sequences of numbers separated by dots, beginning with the letter “A” (e.g., “A12.1.00.001” for “heart muscle”). The first two digits denote an organ (e.g., “A12” for “cardiovascular system”). In this way, each expression value can be uniquely assigned to a tissue or organ, thereby circumventing naming discrepancies when using data from different databases. For each protein, we multiplied the total number of amino acids and the number of selected residues by the protein’s expression value. These weighted values were then summed across all proteins within each tissue and organ to obtain the total amino acid and residue content. We then calculated the overall enrichment of selected amino acids in each tissue and organ by dividing the total content of selected residues by the total amino acid content. For tissues, 13 showed no expression in the HPA transcriptomic dataset, nine in the HPA proteomic dataset, and 18 in the PtDB proteomic dataset. For organs, five showed no expression in the two HPA datasets and four in the PtDB proteomics dataset.

To identify proteins and families with an over-representation of these amino acid groups compared to the rest, we calculated the Z-score, which measures how many standard deviations a value deviates from the mean^18^. This means that a Z-score of one is one standard deviation from the mean, two is two standard deviations, three is three standard deviations (typically defined as outliers), and so on. Proteins and Families were ranked by the Z-scores of their absolute and relative values to emphasise those with particularly high proportions of aromatic heterocycles, non-aromatic heterocycles, carbocycles, and all other amino acids. Spearman rank correlation tests were applied to the tissue and organ datasets as well as the MS dataset to evaluate relationships within a dataset (e.g., protein count vs. enrichment of aromatic heterocycles) and relationships between datasets (e.g., enrichment of aromatic heterocycles in organs vs. intensity of iodine staining)^19^. To ensure the robustness of all tests, we performed permutation tests with 100,000 iterations.

Lastly, each data subset was then analysed separately for the tissue and organ with or without scaling. Analysis of variance (repeated measures ANOVA) was performed to test for significant differences between the perturbation groups (R:aov;^20^). Based on this, all possible groups were compared using a post hoc test (Turkey’s HSD). An additional variance test (Levene) was applied to address the issue of inhomogeneous variance across all parameters. Finally, an additional post hoc test (Games-Howell;^21^) was performed to compare all possible groups when variances were problematic. The statistical analysis was carried out using the freely available software R (version: 4.2.3). The libraries (R.matlab, data.table, Rfast, tidyverse, stats, rstatix, car, ggstatsplot, viridis, openxlsx) were used for specific functions.

## Results

### Data overview and proteins

There were 436,602 aromatic residues overall and 21.5 on average (3.4%), 718,775 non-aromatic heterocycles overall and 35.4 on average (8.7%), 718,401 carbocycles overall and 35.3 on average (14.9%), and 9,509,854 other residues overall and 467.8 on average (73.0%). The overall amino acid amount was 11,383,632 residues with an average of 415 (Fig. 1a).

**Figure 1.**
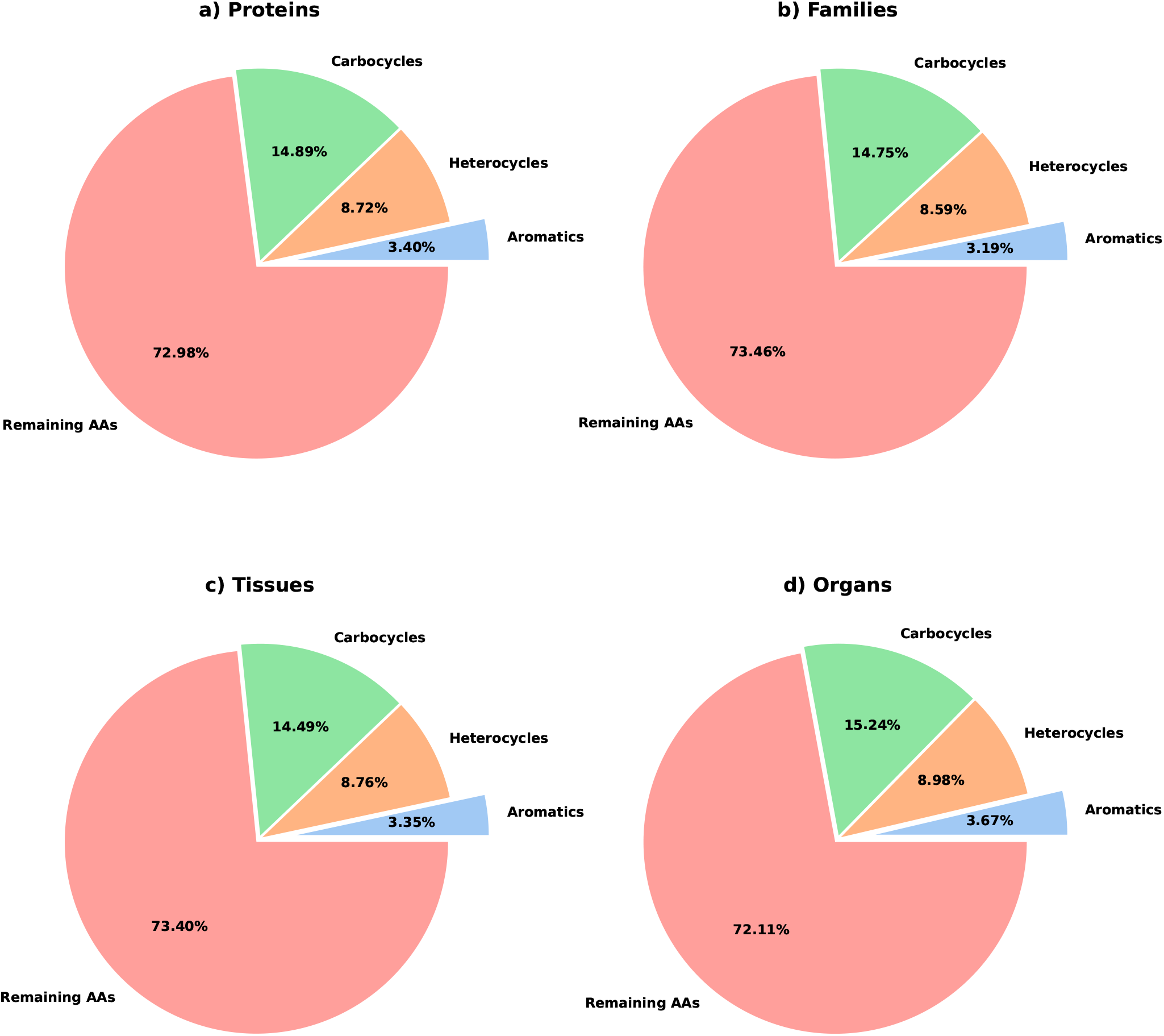
Median enrichment of aromatic heterocycles, non-aromatic heterocycles, carbocycles, and other amino acids in a) proteins, b) families, c) tissues, and d) organs. For tissues and organs, the average of all three expression data sets was taken.

Among proteins with the highest aromatic heterocyclic amino acid content, Titin (*TTN*) stands out with 944 occurrences in its 34,350-residue sequence (*Z* = 40.3), followed by Mucin-3B (*MUC3B*) with 533 in 13,477 residues (*Z* = 22.34) and Filaggrin (*FILA*) with 434 in 4,061 residues (*Z* = 18.0) (Table 1). Naturally, larger proteins tend to have higher absolute counts simply because of their size, although their proportion of heterocyclic amino acids relative to total length remains low. For example, only 2.7% of titin’s residues are heterocyclic, which is below the dataset-wide average of 3.4%.

**Table 1.** The top ten human proteins with the absolute highest amount of aromatic heterocycles, along with their corresponding Z-scores.

| Gene name | Full name | #Aromatic heterocycles | Z-score |
| --- | --- | --- | --- |
| TITIN | Titin | 944 | 40.3 |
| MUC3B | Mucin-3B | 533 | 22.3 |
| FILA | Filaggrin | 434 | 18.0 |
| MUC16 | Mucin-16 | 408 | 16.9 |
| NEBU | Nebulin | 389 | 16.1 |
| SYNE1 | Nesprin-1 | 369 | 15.2 |
| SSPO | SCO-spondin | 314 | 12.8 |
| OBSCN | Obscurin | 298 | 12.1 |
| HORN | Hornerin | 284 | 11.5 |
| DNHD1 | Dynein heavy chain domain-containing protein 1 | 267 | 10.7 |

In contrast, among proteins with an enrichment of heterocyclic amino acid content exceeding 10%, Histidine-rich carboxyl terminus protein 1 (*HRCT1*) has the highest proportion, with 23.4% of its 115 residues being aromatic heterocycles (*Z* = 11.8) (Table 2). It is followed by Putative uncharacterized protein FLJ46204 (*YD019*) and Proline, histidine, and glycine-rich protein 1 (*PHGR1*), with 18.3% (*Z* = 10.3) and 16.0% (*Z* = 8.7) in sequences of 223 and 82 residues, respectively. Since even a few occurrences significantly impact small proteins, they are more likely to exhibit high relative proportions despite their low absolute counts. For example, the Proline, histidine, and glycine-rich protein 1 contains only 15 aromatic heterocyclic amino acids, also below the dataset-wide average of 21.5 residues. The proteins showed a standard deviation of 1.7% for aromatic heterocycles.

**Table 2.** The top ten human proteins with the relatively highest amount of aromatic heterocycles, along with their corresponding Z-scores.

| Gene name | Full name | %Aromatic heterocycles | Z-score |
| --- | --- | --- | --- |
| HRCT1 | Histidine-rich carboxyl terminus protein 1 | 23.5 | 11.79 |
| YD019 | Putative uncharacterized protein FLJ46204 | 21.1 | 10.34 |
| PHGR1 | Proline, histidine and glycine-rich protein 1 | 18.3 | 8.67 |
| PLAC4 | Placenta-specific protein 4 | 16.0 | 7.29 |
| SPGOS | Putative uncharacterized protein SPART-AS1 | 15.4 | 6.92 |
| PRM2 | Protamine-2 | 13.7 | 5.92 |
| HIS3 | Histatin-3 | 13.7 | 5.92 |
| YV018 | Putative uncharacterized protein DKFZp434K191 | 13.7 | 5.92 |
| SNIT1 | Uncharacterized protein encoded by SND1-IT1 | 13.6 | 5.86 |
| S39A7 | Zinc transporter SLC39A7 | 13.4 | 5.74 |

For the MS data, staining values were collected for 11 organs. For the expression data, the HPA transcriptomic dataset contained values for 18,975 proteins, the HPA proteomics dataset for 10,845 proteins, and the PtDB dataset for 16,539 proteins.

### Families

The top families for absolute values showed mostly large, domain-defined families with many members (> 100 proteins), naturally leading to a large amount of aromatic residues (> 4, 000 amino acids) (Fig. 1b, Table 3). The family with the highest amount of aromatic heterocycles is the *Zinc fingers C2H2-type*-family with a total amount of 30,260 residues (*Z* = 30.5) and 763 proteins (the largest group over the entire dataset). This is followed by the *WD repeat domain containing*-family with 10,506 residues (*Z* = 10.4) and the *CD molecules*-family with 7,349 residues (*Z* = 7.2) and 334 proteins (the second largest group). As far as structural protein families are concerned, these are distributed across the entire dataset, mainly families with a low amount of protein content (Table 4). The highest is the *Laminin subunits*-family with 906 residues (*Z* = 0.67) and 12 proteins at rank 59 to as low as rank 1,384 for the *Keratins*-family with eleven residues (*Z* = *−* 0.24) and only one protein.

**Table 3.** The top ten protein families with the absolute highest amount of aromatic heterocycles, along with the number of proteins and their corresponding Z-scores.

| Gene family name | #Proteins | #Aromatic heterocycles | Z-score |
| --- | --- | --- | --- |
| Zinc fingers C2H2-type | 763 | 30,260 | 30.48 |
| WD repeat domain containing | 264 | 10,506 | 10.42 |
| CD molecules | 334 | 7,349 | 7.22 |
| Ring finger proteins | 296 | 7,100 | 6.96 |
| Ankyrin repeat domain containing | 188 | 6,207 | 6.06 |
| MicroRNA protein coding host genes | 256 | 6,069 | 5.92 |
| Pleckstrin homology domain containing | 159 | 5,474 | 5.31 |
| Fibronectin type III domain containing | 94 | 5,297 | 5.13 |
| Armadillo like helical domain containing | 128 | 5,098 | 4.93 |
| EF-hand domain containing | 145 | 4,212 | 4.03 |

**Table 4.** Selected structural protein families with the absolute amount of aromatic heterocycles, along with the number of proteins, their corresponding Z-scores and their rank in the list.

| Gene family name | #Proteins | #Aromatic heterocycles | Z-score | Rank |
| --- | --- | --- | --- | --- |
| Laminin subunits | 12 | 906 | 0.67 | 59 |
| Tubulins | 24 | 395 | 0.15 | 169 |
| Collagens | 8 | 289 | 0.05 | 238 |
| Spectrins | 1 | 188 | -0.06 | 371 |
| Actins | 7 | 90 | -0.15 | 689 |
| Lamins | 3 | 40 | -0.21 | 1,080 |
| Collagens | 8 | 11 | -0.24 | 1,384 |

Protein families with a high enrichment of aromatic heterocycles are dominated by enzymes responsible for membrane and lipid metabolism. These families largely have a low amount of protein content in common (< 10) (Table 5). The highest enrichment is found in the family of *Fatty acid hydroxylase domain containing* with 9.3% (*Z* = 4.9) and six proteins, followed by *Fatty acid desaturases* with 9% (*Z* = 4.6) and eight proteins. In third place, with a Z-score still above 4, is the family of *Alpha-L-fucosidases* with an enrichment of 8.4% (*Z* = 4.1) and two proteins. For structural protein families, a similar pattern to that observed for absolute values can be seen with regard to their enrichment of aromatic heterocycles, namely a distribution across the entire dataset (Table 6). A striking feature here is that the *Spectrins*-family, although only ranked 155th, has an above-average enrichment of 5.1% (*Z* = 1.2). The other structural protein families have Z-scores around zero or below, for example, the *Lamins*-family at rank 1,364 with an enrichment of 2.1% (*Z* = *−* 1.5). The protein families showed a standard deviation of 1.1% for aromatic heterocycles.

**Table 5.**
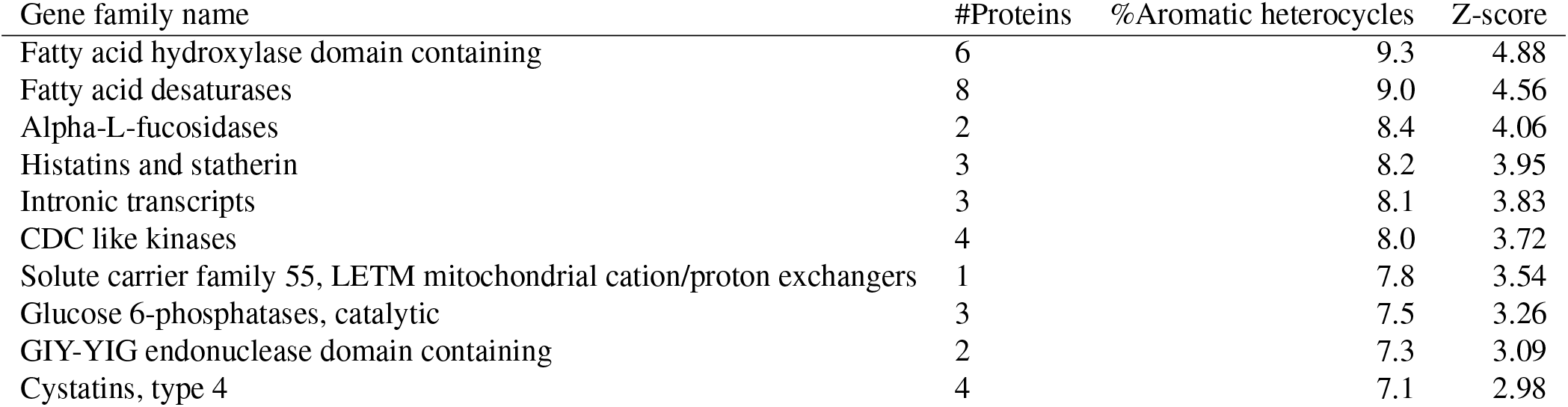
The top ten protein families with the relative highest amount of aromatic heterocycles, along with the number of proteins and their corresponding Z-scores.

**Table 6.** Selected structural protein families with the relative amount of aromatic heterocycles, along with the number of proteins, their corresponding Z-scores and their rank in the list.

| Gene family name | #Proteins | %Aromatic heterocycles | Z-score | Rank |
| --- | --- | --- | --- | --- |
| Spectrins | 1 | 5.1 | 1.82 | 155 |
| Tubulins | 24 | 3.7 | -0.04 | 706 |
| Keratins | 1 | 3.7 | -0.05 | 712 |
| Laminin subunits | 12 | 3.5 | -0.26 | 867 |
| Actins | 7 | 3.4 | -0.32 | 910 |
| Lamins | 3 | 2.1 | -1.46 | 1,364 |
| Collagens | 1 | 2.0 | -1.60 | 1,377 |

### Tissues

In terms of absolute values, tissues with many proteins naturally contain a higher overall number of aromatic heterocycles, as significant correlation coefficients are observed across all three datasets (Fig. 1c). On the other hand, no significant correlations between the three datasets can be identified, except between the two proteomics datasets (*ρ* = 0.49, *p* = 0.0001) (Table 7). Consequently, there are only partial overlaps between the three datasets in the top 20 tissues, mainly between the two proteomics datasets. Only the tissue *Glandula suprarenalis* has a similar rank in all three datasets (HPA transcriptomic: 16, HPA proteomic: 17, PtDB proteomic: 15).

**Table 7.** Spearman rank correlation coefficients of absolute amount of aromatic heterocylces in tissues between the three expression datasets.

| Spearman's $\rho$ | HPA transcriptomic | HPA proteomic | PtDB proteomic |
| --- | --- | --- | --- |
| HPA transcriptomic | 1 |  |  |
| HPA proteomic | 0.23 | 1 |  |
| PtDB proteomic | 0.125 | 0.49 | 1 |

Similar to the absolute values, medium to high correlation coefficients can be observed between the number of proteins in each tissue and their enrichment of aromatic heterocycles. In contrast, the cross-dataset correlations in this case show a low to medium coefficient between the two HPA datasets (*ρ* = 0.37, *p* = 0.005) (Table 8). Consequently, overlaps exist mainly between the two HPA datasets in the top 20 intersections, albeit to a lesser extent. However, it is important to note, especially with regard to the correlation tests, that the standard deviation of the enrichments in all three datasets is very low (HPA transcriptomic: 0.39%, HPA proteomic: 0.21%, PtDB proteomic: 0.11%).

**Table 8.** Spearman rank correlation coefficients of absolute amount of aromatic heterocylces in tissues between the three expression datasets.

| Spearman's $\rho$ | HPA transcriptomic | HPA proteomic | PtDB proteomic |
| --- | --- | --- | --- |
| HPA transcriptomic | 1 |  |  |
| HPA proteomic | 0.365 | 1 |  |
| PtDB proteomic | -0.021 | 0.134 | 1 |

### Organs

As with tissues, there is an almost perfect correlation between the number of proteins and the total amount of aromatic heterocycles in organs, as organs with many proteins naturally express more heterocycles (Fig. 1d). This effect is even more pronounced here than in tissues. In addition, there are high correlations between the three expression datasets, indicating cross-consistency (Table 9). Although high correlations were observed between the datasets, it is difficult to assess overlaps due to the limited number of organs. Furthermore, no significant correlation is observed between the enrichment of aromatic heterocycles and the staining intensity of the respective organs.

**Table 9.** Spearman rank correlation coefficients of absolute amount of aromatic heterocylces in organs between the three expression datasets as well as the MS database.

| Spearman's $\rho$ | HPA transcriptomic | HPA proteomic | PtDB proteomic | MorphoSource |
| --- | --- | --- | --- | --- |
| HPA transcriptomic | 1 |  |  |  |
| HPA proteomic | 0.882 | 1 |  |  |
| PtDB proteomic | 0.758 | 0.869 | 1 |  |
| MorphoSource | 0.242 | 0.107 | 0.022 | 1 |

As with the results for absolute values, a medium to high correlation is returned between the number of proteins in each organ and their enrichment of aromatic heterocycles. Furthermore, similar to the enrichment results in tissues, a significant correlation can be observed between the two HPA datasets (*ρ* = 0.82, *p* = 0.0002) (Table 10). However, due to the low number of organs, it is again difficult to evaluate the overlaps. It is important to note, especially with regard to the correlation tests, that the standard deviation of the enrichments in all three datasets is even lower than in tissues (HPA transcriptomic: 0.32%, HPA proteomic: 0.06%, PtDB proteomic: 0.07%). Moreover, as with absolute values, there are no significant correlations between aromatic heterocycle enrichment and the staining intensity of the respective organs.

**Table 10.** Spearman rank correlation coefficients of relative amount of aromatic heterocylces in organs between the three expression datasets as well as the MS database.

| Spearman's $\rho$ | HPA transcriptomic | HPA proteomic | PtDB proteomic | MorphoSource |
| --- | --- | --- | --- | --- |
| HPA transcriptomic | 1 |  |  |  |
| HPA proteomic | 0.821 | 1 |  |  |
| PtDB proteomic | 0.268 | 0.465 | 1 |  |
| MorphoSource | 0.259 | 0.205 | 0.428 | 1 |

## Discussion

### Proteins with numerous aromatic heterocycles

Of all proteins, titin contains the largest number of aromatic heterocycles. This is unsurprising as it is the largest known human protein with a total amino acid length of ∼30,000. Additionally, it is highly expressed in muscles, as it is a structural protein responsible for their tension^22^. Other important structural proteins in the skeletal musculature are myosin (*MYH2*: 2.3%; *MYH4*: 2.5%; *MYH1*: 2.4%), actin (*ACTS*: 3.4%), nebulin (*NEBU*: 4.6%), and obscurin (*OBSCN*: 3.7%)′^23, 24^. Nevertheless, a relative comparison of titin with these structural proteins shows that they are rich in aromatic heterocycles (2.7%). On the other hand, special isoforms of myosin (*MYH6*: 2.3%; *MYH7*: 2.6%) and actin (*ACTC1*: 3.4%) are found in the heart muscle (Fig. 2). The high proportion of aromatic heterocycles in the structural proteins also explains why the muscles, and in particular the heart muscles, are well stained by iodine^14,25–29^.

**Figure 2.**
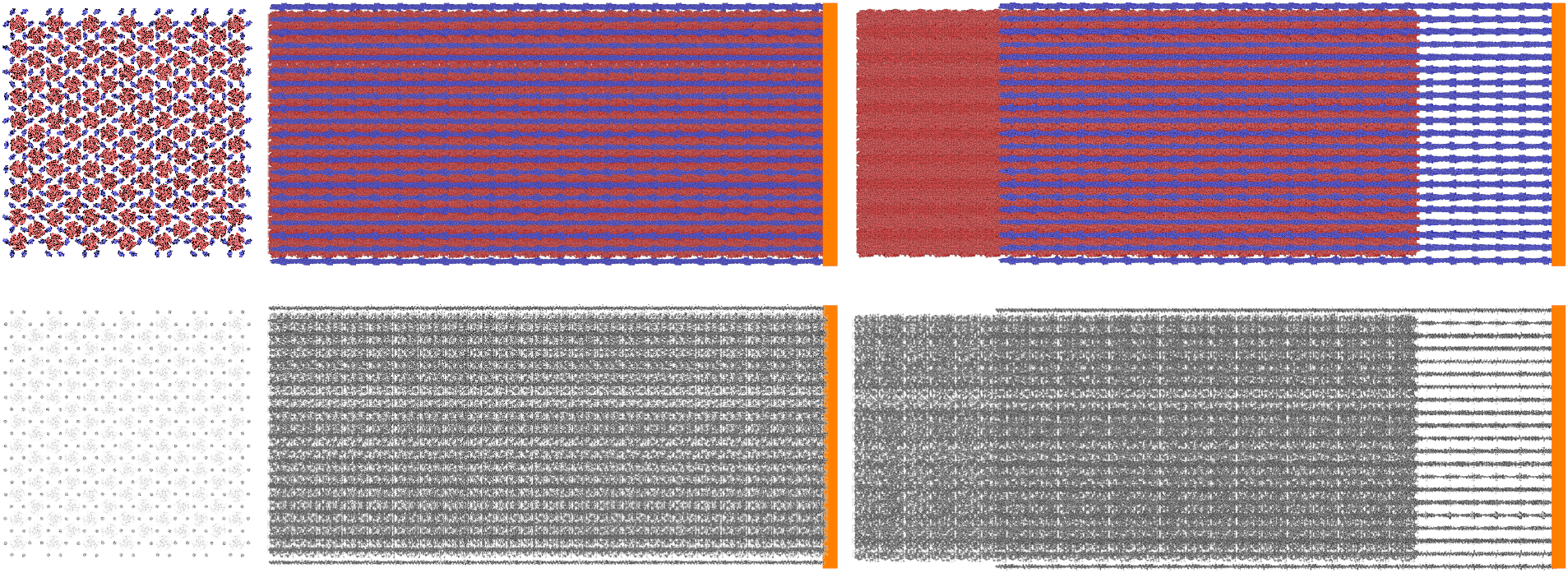
Half-sarcomere protein arrangements of the cardiac muscle (Thin filaments with 7056 protein complexes PDB:6KN7 with a total of 54,902,736 AAs, and thick filaments with 2728 protein complexes PDB:8G4L with a total of 140,322,864 AAs)^40–43^. From left to right: Cross-section at the level of the Z-line (orange), 100% overlap, and 75% overlap of the myosin (red) and actin filament (blue). Below, only the aromatic heterocycles are highlighted in black.

Another structural protein family is the collagens, which are rich in proline but not in aromatic heterocycles^30–32^. Exceptions are collagen types XX (FACITs), XVIII (Endostatin precursor), XXVI (Other), VI (network-forming) and XIV (FACITs) (*COKA1*: 3.5%; *COIA1*: 3.2%; *COQA1*: 2.7%; *CO6A5*: 2.7%; *COEA1*: 2.6%). Almost all type VI collagens (network-forming), which occur in the extracellular matrix, show a high number of aromatic heterocycles (*CO6A1* - *CO6A6*: 1.6% - 2.7%). In contrast, the most frequently occurring collagen types I (fibrillar), IV (network-forming), VIII (network-forming), IX (FACITs) and X (network-forming) contain only a few aromatic heterocycles (*CO1A1*: 1.0%; *CO1A2*: 1.5%; *CO4A5*: 1.1%; *CO8A1*: 1.6%; *COAA1*: 1.8%; *CO8A2*: 1.4%; *CO9A2*: 1.2%), with the exception of *CO4A4* (2.5%). Interestingly, the collagen family sits on close to the bottom of the protein family list at rank 1,384 for absolute value (Table 4) and at rank 1,377 for relative values (Table 6). This could be explained by the fact that the individual proteins are grouped into families, which could reduce the effect of the few highly concentrated collagens. On the other hand, this might explain why areas between the fascicles within the muscle are less intensely stained, allowing the fascicles to be differentiated.

However, structural proteins are not the only ones with high aromatic heterocyclic content. Keratin-associated proteins (max. 6.8%), fillagrin (*FILA*: 10.7% and *FILA2*: 10.6%), and hornerin (*HORN*: 10.0%)^33^. They thus represent a group of iodine-sensitive proteins in the skin. This could explain the high contrast of this tissue when stained with iodine^5^.

Directly downstream in the gastrointestinal tract, the mucins (*MUC1*: 4.7%; *MUC2*: 2.1%; *MUC3A*: 2.5%; *MUC3B*: 4.0%; *MUC4*: 4.0%; *MUC5A*: 2.8%; *MUC5B*: 3.5%; *MUC6*: 4.7%; *MUC7*: 3.2%) with their high number of aromatic heterocycles represent another important group. Mucins are structural components of mucus on mucous membranes, including the gastrointestinal tract. We know from Fennerty (1999) that the oesophagus can be easily stained with iodine/Lugol’s solution. This suggests that mucins play a significant role in this process. Furthermore, increased mucin production is observed in many types of cancer^34,35^. *PHGR1* (18.3%), another gene expressed in the gastrointestinal tract, tops the relative proportions of aromatic heterocycles.

The best-known reaction involving iodine occurs in the thyroid gland^36^. However, this is an enzymatic reaction and is based on the production of iodine-containing thyroid hormones, such as triiodothyronine (T3) and thyroxine or tetraiodothyronine (T4), and the peptide hormone calcitonin. The bound iodine levels are negligible (1.17 ng/dL), as these hormones occur at very low concentrations compared to those of the aromatic heterocycles.

### Families, tissues, and organs

To analyse the large amount of data, we grouped the proteins into four levels of complexity: individual proteins, protein families, tissues, and organs. Organs were especially interesting as we could directly compare their enrichments of aromatic heterocycles to iodine staining intensity, which was not available for families and tissues. In general, the higher the complexity level (protein -> family -> tissue -> organ), the less differentiable the ingroups are, as shown by the variability in their enrichments (protein: 1.7%, family: 1.1%, tissue: (max.) 0.39%, organ: (max.) 0.32%).

As mentioned in the results, enzyme families rank among the highest for aromatic heterocyclic enrichment. However, there are far fewer enzyme proteins than structural proteins in tissues^37^. The only structural protein family that is notable is the *Spectrins*-family with an above-average enrichment of 5.1% at rank 155 (*Z* = 1.82). In this context, it is important to note that there can be large variations in enrichments within a family, which can distort the data and, consequently, lead to a low ranking.

As already mentioned, we were unable to find any correlation between the absolute number or enrichment of aromatic heterocycles in organs and their iodine staining intensity. One of the main reasons for this is that organs consist of many proteins and tissues, which leads to a high degree of protein aggregation. This can be explained using the example of muscle: it contains proteins with a high affinity for iodine (e.g., myosin, actin, titin), whereas other proteins do not have this affinity (e.g., collagens). In addition, the grouping in the three expression datasets is insufficient. For example, the two HPA datasets contain only one tissue assigned to muscle, with a low number of proteins (HPA transcriptomics: 15,974; HPA proteomics: 5,811), whereas the PtDB dataset shows no expression in this organ. This also applies to other tissues and organs (e.g., “cardiovascular system”).

## Conclusions

In conclusion, iodine binds to proteins to varying extents, enabling the targeted staining and classification of specific tissues. Importantly, it is primarily the overall protein content within the tissue that determines staining intensity, rather than the aromatic heterocyclic content of individual proteins. For example, tissues or organs with low protein content are difficult to stain, such as fatty, bony and water-rich tissues.

## Outlook

The method presented here is particularly well-suited for application to other organisms, offering a targeted approach for staining specific tissues. Future studies could also explore how iodination affects protein stability, as was done earlier for various types of non-enzymatic protein modification, such as bromination and chlorination^38^. Susceptibility scores can be determined for particular amino acids and then combined into overall scores for an entire protein. Notably, many top protein hits are localised in the skin and digestive tract, tissues frequently exposed to high microbial loads. Given that iodine, particularly in the form of Lugol’s solution, is known for its antimicrobial properties^39^, this observation provides a compelling basis for further investigation.

## Data availability

The data supporting this study’s findings are available at DOI: 10.17632/nd4ch2vhty

## Acknowledgements

We would like to thank Rainer Beckert, all members of the Department of Bioinformatics (especially Prof. Schuster), and the Institute of Zoology and Evolutionary Research. H.S. was supported by DFG FI 410/24-1.

## Author contributions statement

H.S., V.W. and L.M.B. designed the study. V.W. performed the statistics. V.W. and H.S. drafted the manuscript and all figures. All authors contributed to the interpretation of the results and revised the manuscript.

